# HIV-2/SIV Vpx antagonises NF-*κ*B activation by targeting p65

**DOI:** 10.1101/2021.06.26.450035

**Authors:** Douglas L. Fink, James Cai, Matthew V. X. Whelan, Christopher Monit, Carlos Maluquer de Motes, Greg J. Towers, Rebecca P. Sumner

## Abstract

The NF-*κ*B family of transcription factors and associated signalling pathways are abundant and ubiquitous in human immune responses. Activation of NF-*κ*B transcription factors by viral pathogen-associated molecular patterns, such as viral RNA and DNA, is fundamental to anti-viral innate immune defences and pro-inflammatory cytokine production that steers adaptive immune responses. Diverse non-viral stimuli, such as lipopolysaccharide and cytokines, also activate NF-*κ*B and the same anti-pathogen gene networks. Viruses adapted to human cells often encode multiple proteins aimed at varied NF-*κ*B pathway targeted to mitigate the anti-viral effects of NF-*κ*B-dependent host immunity. In this study we have demonstrated using numerous assays, in a number of different cell types, that plasmid-encoded or virus-delivered Simian Immunodeficiency Virus (SIV) accessory protein Vpx is a broad antagonist of NF-*κ*B signalling active against diverse innate NF-*κ*B agonists. Using targeted Vpx mutagenesis, we showed that this novel Vpx phenotype is independent of known Vpx cofactor DCAF1 and other cellular binding partners, including SAMHD1, STING and the HUSH complex. We found that Vpx co-immunoprecipitated with canonical NF-*κ*B transcription factor p65 and not NF-*κ*B transcription factor proteins p50 or p100, preventing nuclear translocation of p65, a novel mechanism of NF-*κ*B antagonism by lentiviruses. We found that broad antagonism of NF-*κ*B activation by Vpx was conserved across distantly related lentiviruses as well as for Vpr from SIV Mona monkey (SIVmon), which has Vpx-like SAMHD1-degradation activity.

**Importance:** Broad antagonism of NF-*κ*B activation has been described for HIV-1, but not for Vpx-encoding lentiviruses such as HIV-2. Here we extend our understanding of lentiviral antagonism by identifying an interaction between Vpx and transcription factor NF-*κ*B p65, leading to inhibition of its nuclear translocation and broad NF-*κ*B antagonism. This further evidences a requirement for lentiviruses to target universal regulators of immunity, including NF-*κ*B, to avoid the anti-viral sequelae of pro-inflammatory gene expression stimulated by both viral and extra-viral agonists, such as lipopolysaccharide translocation, during disruption of the gut microbiome barrier during HIV-1 infection. Further structural studies of p65 targeting by Vpx may yield translational insights in the form of novel pan-NF-*κ*B inhibitors for pathologies characterised by excessive NF-*κ*B activity. Our findings are also relevant to the gene therapy field where virus-like particle associated Vpx is routinely used to enhance vector transduction through antagonism of SAMHD1, and perhaps also through manipulation of other pathways such as NF-*κ*B.

## Introduction

NF-*κ*B transcription factors (TFs) are pivotal mediators of inflammation(1). Amongst diverse roles in immune signalling, NF-*κ*B activation is fundamental to pro-inflammatory anti-viral innate defences. Viruses typically activate NF-*κ*B through detection of viral pathogen-associated molecular patterns (PAMPs) by host pattern-recognition receptors (PRRs), such as Toll-like receptors, retinoic acid-inducible gene (RIG)-like receptors and DNA sensors (such as the cyclic GMP-AMP synthase (cGAS) and stimulator of interferon genes (STING) pathway)(2–5). Canonical NF-*κ*B activation by PRRs, and other diverse ligands such as cytokines, leads to degradation of inhibitory proteins (such as IκB) and release of NF-*κ*B TF dimers, predominantly p50 and p65 (RelA), which translocate into the nucleus to upregulate an array of genes that govern cell-autonomous and adaptive responses against infection(1,6,7). Non-canonical NF-*κ*B pathways are activated by a more narrow range of stimuli, including a subset of the TNF receptor superfamily members such as CD40, and depend on processing of TF p52 precursor protein, p100(8).

Unsurprisingly, viruses have evolved strategies to antagonise NF-*κ*B signalling, often targeting pathway activation at multiple levels in complex ways, emphasising the primacy of NF-*κ*B TFs to host defences(9). HIV-1 deploys at least two direct NF-*κ*B antagonists which target multiple points of NF-*κ*B signalling. Accessory protein Vpu, encoded by HIV-1 and its ancestor lentivirus, chimpanzee simian immunodeficiency virus (SIVcpz), antagonises NF-*κ*B through at least two mechanisms. Vpu directly binds and inhibits tetherin, a host restriction factor that inhibits lentiviral budding and also activates NF-*κ*B on virus engagement (10–12). Independently of tetherin, Vpu also stabilises IκB to inhibit p65 nuclear translocation and NF-*κ*B-activated transcription(13,14). In contrast, HIV-2 does not possess a *vpu* gene and tetherin antagonism is effected through HIV-2 Env and/or Nef (Ref). Intriguingly Env activates rather than inhibits NF-*κ*B signalling(15). HIV-1 also suppresses NF-*κ*B activation at the level of TF nuclear transport, through accessory protein HIV-1 Vpr interaction with karyopherins(16,17). This equivalent activity of HIV-1 Vpu or Vpr as broad NF-*κ*B antagonists has not been studied for HIV-2.

HIV-2-related SIVsm/SIVmac and SIVrcm/SIVmnd-2 lineage lentiviruses, which infect sooty mangabey, macaque, red-capped mangabey and mandrill monkeys respectively, all of which belong to the same primate family, possess two homologous genes to HIV-1/SIVcpz *vpr. vpr* and *vpx*(18,19). Typically if a virus has 2 Vpr-like genes, one of them is named Vpx. That is, no viruses have been assigned a Vpx without a Vpr. But the identification of a particular gene as a Vpr or a Vpx is complex because high levels of adaptation prevent alignment and effective phylogenetic analysis and functions overlap between the two proteins(20). Like Vpr, Vpx is packaged into lentiviral virions consistent with its involvement in early events of the lentiviral lifecycle counteracting host innate defences(21). Like Vpr, Vpx interacts with host interactor protein damage-specific DNA binding protein 1 (DDB1)-Cullin4A (CUL4A)-associated factor 1 (DCAF1) which promotes ubiquitination and drives recruitment of proteasome machinery to degrade target host proteins, most notably sterile alpha motif and histidine-aspartate domain containing protein 1 (SAMHD1) and the human silencing hub (HUSH) complex, in order to enhance virus replication(22–24).

Here we demonstrate that Vpx is a broad inhibitor of NF-*κ*B activation and pro-inflammatory gene expression active against diverse NF-*κ*B agonists, including during virus infection. Vpx recruits p65 and inhibits nuclear translocation independently of Vpx-cofactor DCAF1. We found that this novel DCAF1-independent phenotype is conserved for all Vpx-encoding lentiviruses tested. We propose that Vpx has evolved to suppress inflammatory signals from a broad range of inflammatory and/or defensive stimuli which would otherwise limit transmission and ongoing replication of Vpx bearing viruses (25).

## Results

### Vpx is a broad antagonist of NF-*κ*B activation

Whilst investigating innate immune responses to SIVsm lineage viruses in primary human immune cells we noted that although wild-type SIVsm infection of human monocyte-derived macrophages (MDM) did not induce NF-*κ*B-dependent gene expression at the doses tested, basal expression of NF-*κ*B-dependent genes such as tumour necrosis factor (TNF)α (Fig 1A) and IL-8 (Fig 1B) were significantly reduced in SIVsm-infected, compared to mock-infected, cells. This inhibition was not observed during infection with a virus lacking Vpx (SIVsmΔvpx) (Figs 1A, B), despite similar infection levels (Suppl. Fig. 1A). We also found that Vpx delivered by genome-free virus-like particles (VLPs) antagonised NF-*κ*B-dependent gene expression activated by lipopolysaccharide (LPS) treatment of MDM (Fig 1C), indicating that Vpx could antagonise NF-*κ*B activation driven by exogenous non-viral agonists. To study this inhibitory activity further we turned to reporter gene assays in HEK293T cells transiently expressing Vpx and an NF-*κ*B-sensitive luciferase construct. Using this system we found that Vpx antagonised NF-*κ*B activation in response to a broad range of stimuli including cytokines TNFα (Fig 1D) and interleukin 1β (IL-1β, Fig 1E), activation of RNA sensing pathways by Sendai virus (SeV, Fig 1F) and activation of DNA sensing by transient over-expression of cGAS and STING (Fig 1G). Inhibition of NF-*κ*B downstream of cGAS/STING was further confirmed by measuring transcripts for NF-*κ*B-dependent gene *CXCL-10* by qRT-PCR (Fig 1H). Antagonism of DNA sensing-induced NF-*κ*B activation by Vpx was dose-dependent (Suppl. Fig 1B) and was not due to inhibition of cGAS or STING expression (Suppl. Fig 1C, D) or cGAS/STING degradation (Suppl. Fig. 1B). In contrast to the effect of Vpx on NF-*κ*B-dependent gene expression, transient co-expression of Vpx with cGAS/STING slightly increased activation of an IRF-3-driven luciferase reporter bearing the *IFIT-1* promoter (also known as *ISG56*, Suppl. Fig 1E, F), as well as *IFIT-1* mRNA expression (Suppl. Fig 1G). Finally, we demonstrated that infection of HEK293T cells with SIVsm inhibited cGAS/STING-dependent NF-*κ*B reporter activation, whilst infection with SIVsmΔvpx did not (Fig 1I), despite equivalent levels of infection (Suppl. Fig 1H). Together these data demonstrated Vpx to be a broad antagonist of NF-*κ*B activated by both cognate virus infection and by exogenous stimuli. Further, in the case of DNA sensing, Vpx specifically inhibited NF-*κ*B activation without affecting the activation of IRF3, demonstrating the specificity of this activity.

**Figure 1.**
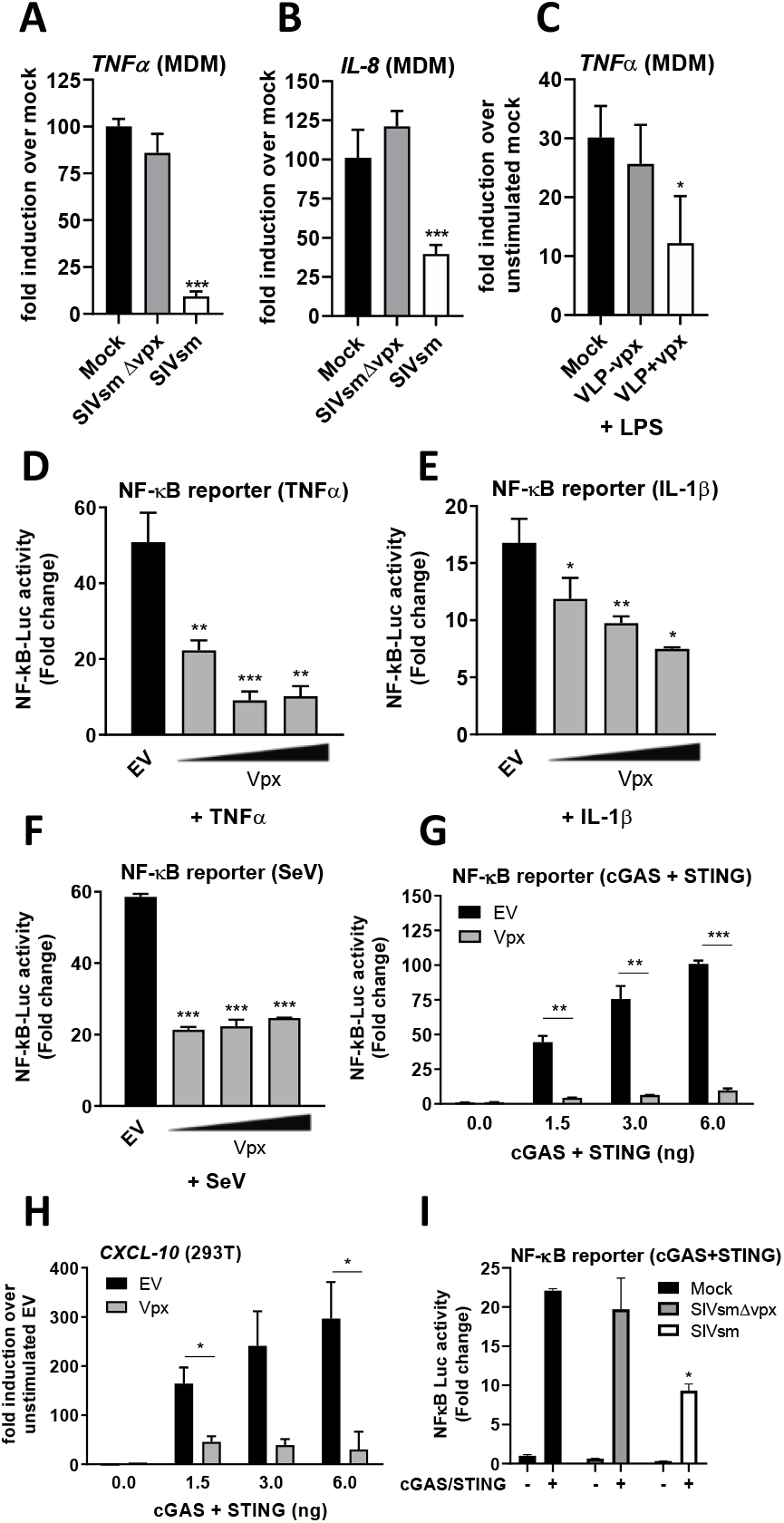
Vpx is a broad antagonist of NF-*κ*B. A: *TNF-α* qRT-PCR from primary human monocyte-derived macrophages (MDM) infected for 48 h with SIVsm or SIVsmΔVpx (1.5 U/ml RT). B: *IL-8* qRT-PCR from MDM infected for 48 h with SIVsm or SIVsmΔVpx (1.5 U/ml RT). C: *TNF-α* qRT-PCR from MDM transduced for 24 h with VLPs -/+ Vpx (3 U/ml RT) followed by stimulation with 1 ng/ml LPS for 24 h. D: NF-*κ*B reporter activity from HEK293T cells transfected for 24h with 25, 50 or 100ng SIVmac Vpx or EV control (100ng) per well and then stimulated for 8 h with 10 ng/ml TNF-α. E: NF-*κ*B reporter activity from HEK293T cells transfected for 24h with 25, 50 or 100ng SIVmac Vpx or EV control (100ng) per well and then stimulated for 8 h with 1 ng/ml IL-1 β. F: NF-*κ*B reporter activity from HEK293T cells transfected for 24h with 25, 50 or 100ng SIVmac Vpx or EV control (100ng) per well and then stimulated for 12 h with 2.0 HA U/ml Sendai virus (SeV). G: NF-*κ*B reporter activity from HEK293T cells co-transfected for 24h with 50ng SIVmac Vpx or EV control plus 0, 1.5, 3 or 6ng each of FLAG-cGAS and FLAG-STING per well. H: *CXCL-10* qRT-PCR from HEK293T cells co-transfected for 24h with 50ng SIVmac Vpx or EV control plus 0, 1.5, 3 or 6ng each of FLAG-cGAS and FLAG-STING per well. I: NF-*κ*B reporter activity from HEK293T cells transfected for 24h with 1.5ng FLAG-cGAS and FLAG-STING per well and then infected for 24 h with SIVsm or SIVsmΔVpx (1.0 U/ml RT). Data are mean ± SD, *n* = 3, representative of at least 3 repeats. Statistical analyses were performed using Student’s *t*-test, with Welch’s correction where appropriate. \**P* < 0.05, \*\**P* < 0.01, \*\*\**P* < 0.001.

### Vpx inhibits NF-*κ*B downstream of p65

To gain mechanistic insight into Vpx antagonism of NF-*κ*B, we undertook pathway mapping in HEK293T cells transiently expressing an NF-*κ*B-sensitive luciferase reporter (Fig 2A). Consistent with the ability of Vpx to inhibit NF-*κ*B activation downstream of diverse stimuli, Vpx inhibited NF-*κ*B activation by expression of adaptor proteins TNFα-activated tumour necrosis factor receptor-associated factor 2 (TRAF2; Fig 2B), and IL-1β and Toll-like receptor (TLR)-activated TRAF6 (Fig 2C), the kinase inhibitor of *κ*B (I*κ*B) kinase β (IKKβ; Fig 2D) and also downstream of exogenous p65 expression (Fig 2E). Together these data suggested antagonism of NF-*κ*B signalling at or after p65 subunit activation.

**Figure 2.**
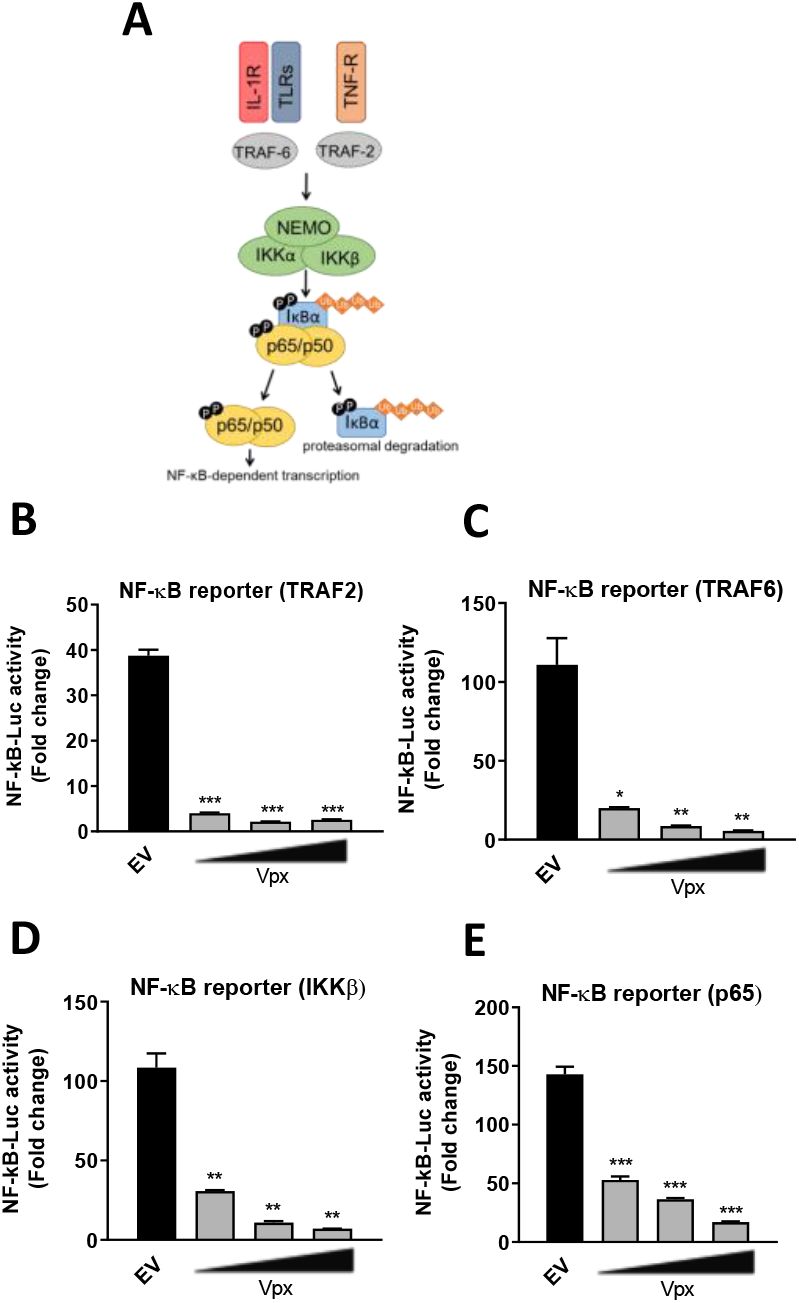
Vpx inhibits NF-*κ*B downstream of p65. A: Schematic of NF-*κ*B activation downstream of the TNF receptor (TNFR), IL-1 receptor (IL-1R) and Toll-like receptors (TLRs). B: NF-*κ*B reporter activity from HEK293T cells co-transfected for 24h with 25-100ng SIVmac Vpx or EV control (100ng) and 25ng of TRAF2. C: NF-*κ*B reporter activity from HEK293T cells co-transfected for 24h with 25-100ng SIVmac Vpx or EV control (100ng) and 25ng of TRAF6. D: NF-*κ*B reporter activity from HEK293T cells co-transfected for 24h with 25-100ng SIVmac Vpx or EV control (100ng) and 50ng of IKKβ. E: NF-*κ*B reporter activity from HEK293T cells co-transfected for 24h with 25-100ng SIVmac?? Vpx or EV control (100ng) and 25ng of p65. Data are mean ± SD, *n* = 3, representative of at least 4 repeats. Statistical analyses were performed using Student’s *t*-test, with Welch’s correction where appropriate. \**P* < 0.05, \*\**P* < 0.01, \*\*\**P* < 0.001.

### Vpx inhibition is independent of DCAF1, SAMHD1 and HUSH

Vpx interacts with a variety of host cellular proteins(26). We used mutagenesis and depletion experiments to determine if these binding partners contributed to NF-*κ*B inhibition. Vpx mutants deficient for antagonism of SAMHD1 (E15A E16A) and HUSH complex (Q47A V48A) retained capacity to antagonise gene expression activated by cGAS/STING (Fig 3A) or p65 expression (Fig 3B)(23,27). A mutant of Vpx that was recently described to prevent interaction with STING (R51AS52A) also still antagonised gene expression activated by p65 expression in a dose-dependent manner as effectively as wild-type Vpx (Fig 3C). This was also true for Vpx Q76R, which is deficient for DCAF1 binding, suggesting DCAF1 independence for this activity (Fig3D)(22,27,28). Concordantly, Vpx inhibited NF-*κ*B activation in HEK293T cells depleted of DCAF1 by siRNA (Figs 3E, F). All mutant Vpx proteins were expressed at similar levels to wild-type (Suppl. Figs 2A, B). Interestingly depletion of DCAF1 itself consistently reduced NF-*κ*B reporter activity downstream of p65 over-expression in the empty vector (EV) control, but inhibition of the remaining signal was still observed by Vpx and this was to the same degree as in siCtrl cells (see normalised data in Fig 3F). Depletion of DCAF1 protein to undetectable levels was confirmed by immunoblotting (Suppl. Fig 3C). These data suggest that Vpx antagonism of NF-*κ*B signalling is independent of DCAF1 and associated Vpx interactome.

**Figure 3.**
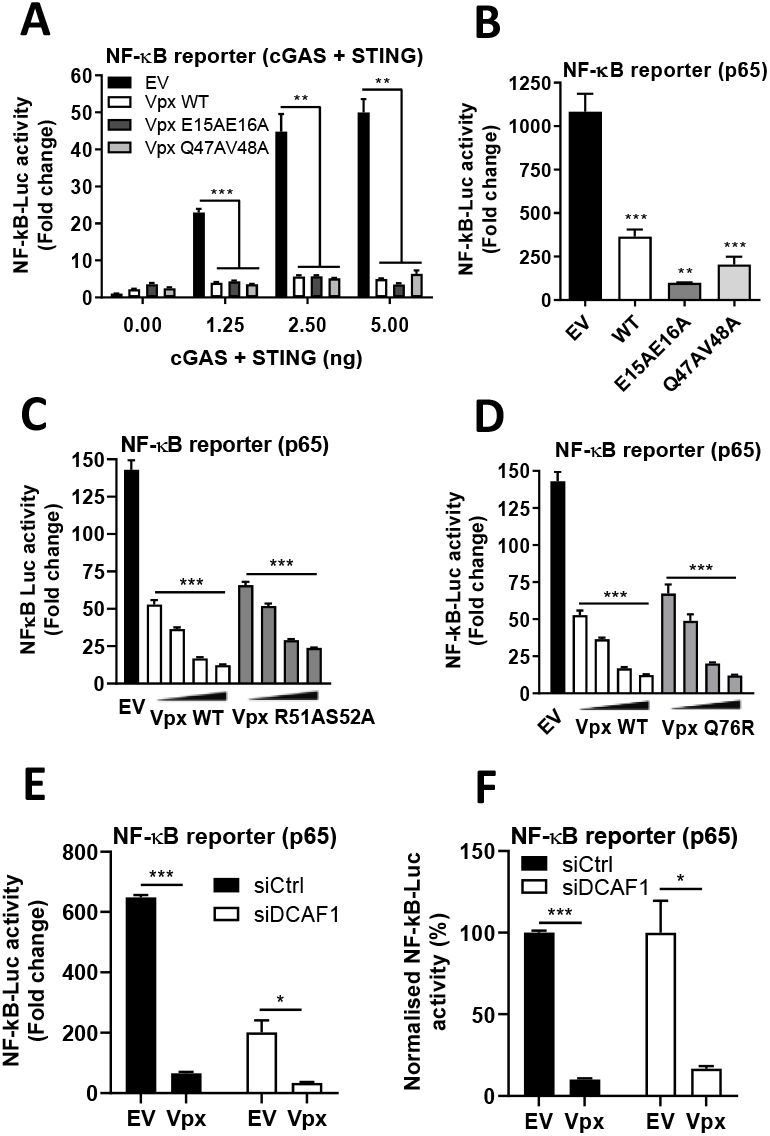
Vpx-mediated inhibition of NF-*κ*B is independent of DCAF1, SAMHD1 and HUSH. A: NF-*κ*B reporter activity from HEK293T cells co-transfected for 24h with 50ng SIVmac; WT, E15AE16A SAMHD1 mutant or Q47AV48A HUSH mutant Vpx or EV control plus 0, 1.2>5, 2.5 or 5ng each of FLAG-cGAS and FLAG-STING per well. B: NF-*κ*B reporter activity from HEK293T cells co-transfected for 24h with 50ng SIVmac WT, E15AE16A SAMHD1 mutant or Q47AV48A HUSH mutant Vpx or EV control plus 25ng p65 per well. C: NF-*κ*B reporter activity from HEK293T cells co-transfected for 24h with 25, 50 or 100 ng SIVmac WT or R51AS52A STING mutant Vpx or EV control plus 25ng p65 per well. D: NF-*κ*B reporter activity from HEK293T cells co-transfected for 24h with 25, 50 or 100 ng SIVmac WT or Q76R DCAF1 mutant Vpx or EV control plus 25ng p65 per well. E: NF-*κ*B reporter activity from HEK293T cells previously transfected for 48h with siRNA against DCAF1 (siDCAF1) or Ctrl siRNA (siCtrl) and then co-transfected for 24h with 50ng SIVmac or EV control plus 25ng p65 per well. F: Normalised NF-*κ*B reporter activity from 3E. Data are mean ± SD, *n* = 3, representative of at least 3 repeats. Statistical analyses were performed using Student’s *t*-test, with Welch’s correction where appropriate. \**P*<0.05, \*\**P* < 0.01, \*\*\**P* < 0.001.

### Vpx interacts with p65

Given that Vpx could inhibit NF-*κ*B induced gene expression by p65 over-expression we tested whether Vpx might bind NF-*κ*B p65 directly. We co-expressed HA-tagged NF-*κ*B proteins from both class I (p100 and p50) and II (p65) as well as IKKβ with FLAG-tagged Vpx in HEK293T cells. Indeed, immunoprecipitation of HA-tagged p65 specifically co-immunoprecipitated Vpx (Fig 4A). This result was confirmed by reciprocal co-immunoprecipitation of HA-p65 on immunoprecipitation of FLAG-Vpx with anti-FLAG antibody (Fig 4B).

**Figure 4.**
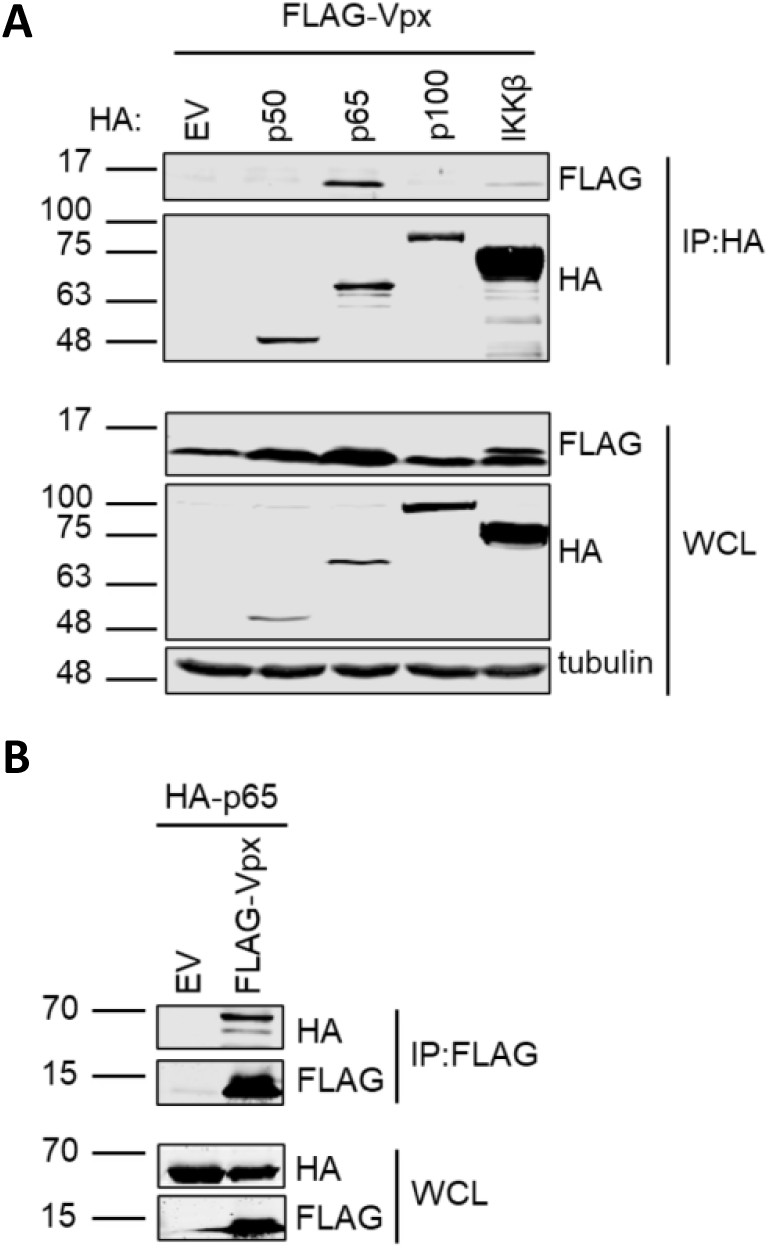
Vpx interacts with p65. A: Immunoblot from co-immunoprecipitation assay performed from HEK293T cells co-transfected with FLAG-Vpx and HA-tagged p50, p65, p100, IKKβ or an EV control. Whole cell lysates (WCL) were probed with FLAG, HA and tubulin antibodies and immunoprecipitates (IP) were probed for FLAG and HA following immunoprecipitation with anti-HA beads. B: Immunoblot from co-immunoprecipitation assay performed from HEK293T cells co-transfected with HA-p65 and FLAG-Vpx or an EV control. WCL and IP samples that had been incubated with anti-FLAG beads were probed with HA and FLAG antibodies. Data are from a representative experiment repeated 4 times.

### Vpx blocks p65 nuclear translocation

To further characterise the impact of Vpx binding p65 we performed assays measuring p65 phosphorylation and nuclear translocation. Phosphorylation of p65 at serine 536 was readily observed in HEK293T cells stimulated with TNFα at 15 and 30 minutes post-stimulation Vpx-transfected cells, indicating that interaction of Vpx with p65 did not interfere with this stimulation-induced phosphorylation event (Fig 5A). Total p65 levels were also unaffected by Vpx expression (Fig 5A), further supporting a model in which inhibition of p65 by Vpx is independent of DCAF1 and concordantly non-degradative (Fig 3D-F). Furthermore, other hallmarks of NF-*κ*B activation such as IκBα phosphorylation and degradation were also unaffected by Vpx expression (Fig 2). Conversely, as a control, vaccinia virus protein B14, an inhibitor of the IKK complex(29), prevented degradation of I*κ*Bα and also reduced phosphorylation of p65, particularly at 30 minutes post-TNFα treatment (Fig 5A). NF-*κ*B inhibition by Vpx and B14 was demonstrated in a parallel reporter gene assay performed using the same conditions as the phospho-blot assay (Fig 5B). Despite phosphorylation of p65 at serine 536 in the presence of Vpx, nuclear translocation of this transcription factor was blocked by Vpx in TNFα-treated cells (Fig 5C, D) explaining inhibition of NF-*κ*B-dependent transcription by inhibition of NF-*κ*B nuclear transport.

**Figure 5.**
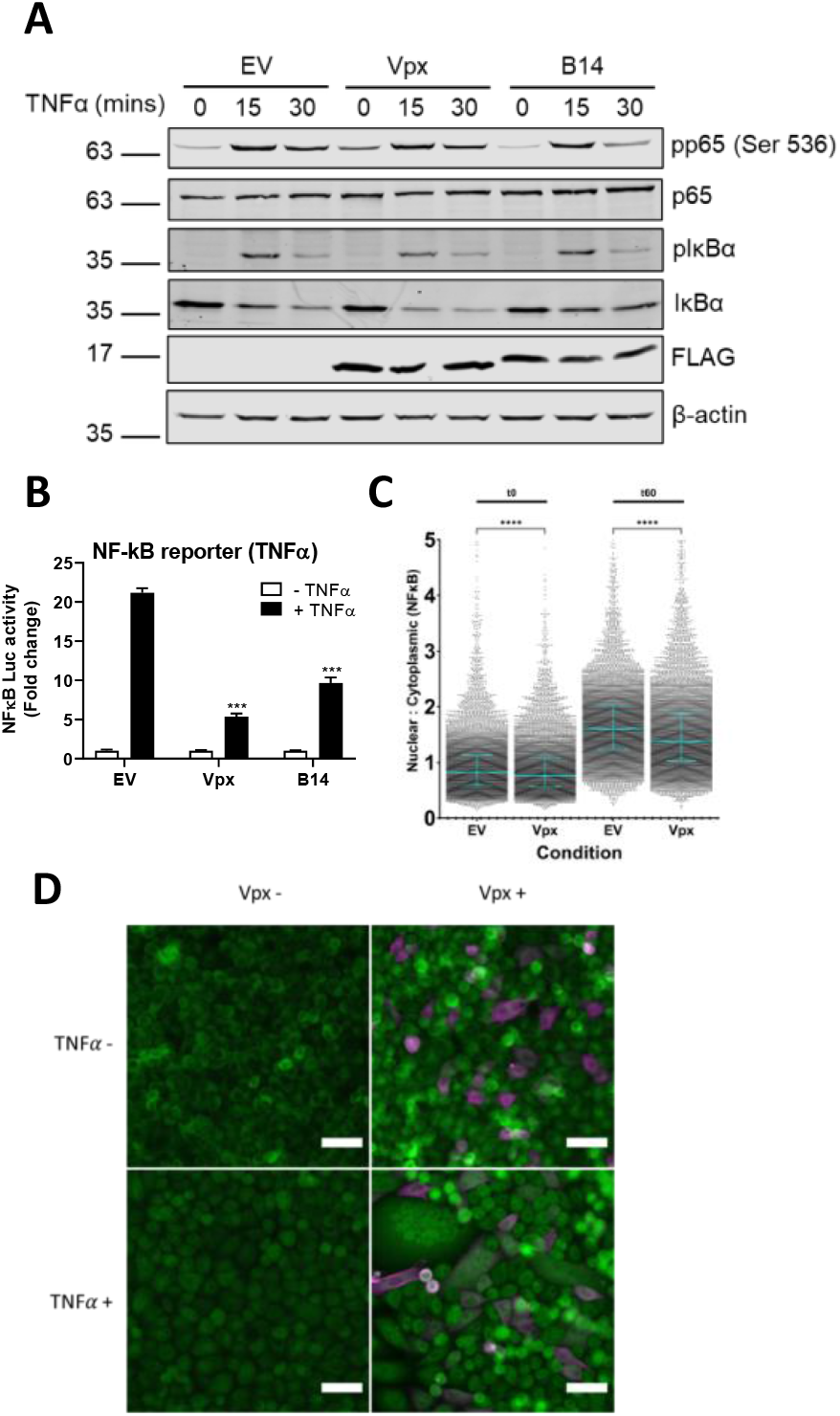
Vpx blocks p65 nuclear translocation. A: Immunoblot from HEK293T cells that had been transfected for 24h with 2μg FLAG-tagged Vpx, vaccinia virus protein B14 or EV control and stimulated for 0, 15 or 30 mins with 50ng/ml TNFα. Blots were probed with antibodies against total p65, phosphorylation of p65 on serine 536 (pp65), total IκBα, phosphorylated IκBα (pIκBα), FLAG for Vpx and B14 expression and actin. B: NF-*κ*B reporter activity from HEK293T cells transfected in parallel with the experiment from 5A and stimulated with 50ng/ml TNFα for 8h. C: Single cell analysis quantifying the Integrated Nuclear Intensity of NF-*κ*B p65 in HeLa cells transfected with FLAG-tagged Vpx or EV control and stimulated for 0 and 30 minutes with 50ng/ml TNFα. Horizontal lines indicate the mean. Kruskall-Wallis test with Dunn’s multiple comparison, ****, *P* <0.0001. D: Representative example of immunofluorescence staining of NF-*κ*B p65 (green) after FLAG-tagged Vpx or EV control transfection and stimulated for 0 and 30 minutes with 50ng/ml TNFα. FLAG-tagged Vpx (magenta). Data in 5A are from a representative experiment repeated 3 times. Data in 5B are mean ± SD, *n* = 3, representative of 3 repeats. Statistical analyses were performed using Student’s *t*-test, with Welch’s correction where appropriate. \*\*\**P* < 0.001.

### Inhibition of NF-*κ*B is conserved amongst Vpx species variants

Vpr/Vpx proteins capable of degrading SAMHD1 are encoded by diverse primate lentiviruses including HIV-2 (Suppl. Fig 6A) (30). To determine whether inhibition of NF-*κ*B was a conserved Vpx feature we cloned a series of diverse Vpx variants and tested their ability to suppress p65-driven NF-*κ*B reporter gene activation in HEK293T cells. In agreement with data obtained using SIVsm infection (Fig 1A, B and I), expression of Vpx from SIVsm strain E543 and HIV-2 inhibited NF-*κ*B reporter activity similarly to Vpx from SIVmac (Fig 6A). The other major clade of lentiviruses encoding genes commonly referred to as Vpx, besides the SIVsm/HIV-2 lineage, are derived from Red Capped Mangabeys and Mandrills (Supplementary Fig 6A) and Vpx from these species also demonstrated anti-NF-*κ*B reporter activity, despite difficulties in detecting SIVrcm Vpx expression by immunoblot (Fig 6B). Finally we tested the Vpr proteins with SAMHD1-degrading activity from lentiviruses infecting vervet (SIVagm_VER) and mona monkeys (SIVmon) and found that whilst SIVagm_VER Vpr did not inhibit NF-*κ*B downstream of p65, SIVmon Vpr did (Fig 6C). Both Vpr proteins expressed well in this assay. Overall this suggests that inhibition of NF-*κ*B is a conserved feature of Vpx, mapping downstream of p65 activation.

**Figure 6.**
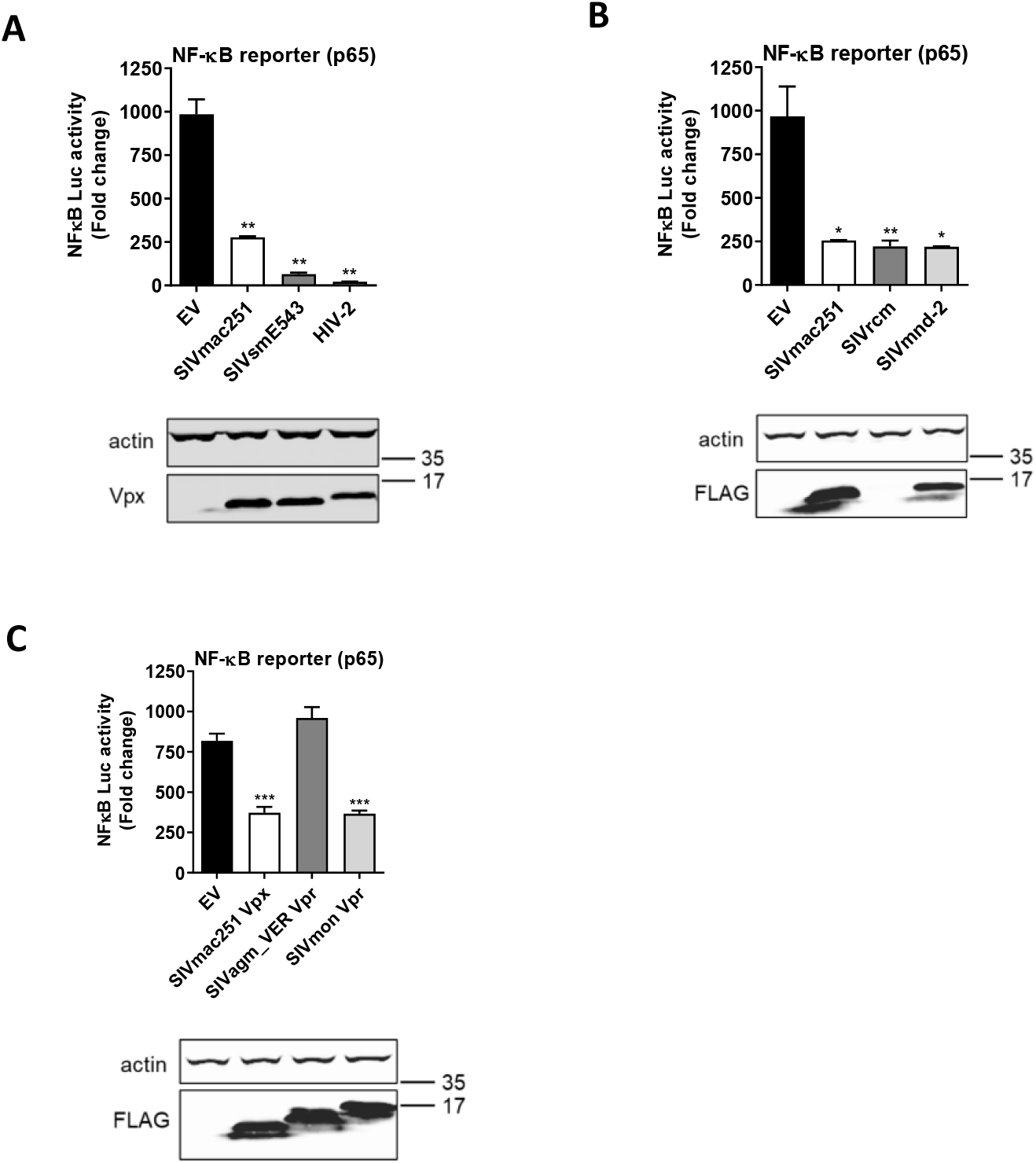
Inhibition of NF-*κ*B is conserved amongst Vpx species variants. A: NF-*κ*B reporter activity and immunoblot from HEK293T cells co-transfected for *24h* with 50ng Vpx proteins from SIVmac strain 251, SIVsm strain E543 or HIV-2 or EV control and 50ng p65 per well. Vpx expression was detected using an anti-Vpx antibody. B: NF-*κ*B reporter activity and immunoblot from HEK293T cells co-transfected for 24h with 50ng FLAG-tagged Vpx proteins from SIVmac strain 251, SIVrcm or SIVmnd-2 or EV control and 50ng p65 per well. Vpx expression was detected using an anti-FLAG antibody. C: NF-*κ*B reporter activity and immunoblot from HEK293T cells co-transfected for 24h with 50ng FLAG-tagged Vpx protein from SIVmac strain 251 or FLAG-tagged Vpr proteins from SIVagm_VER or SIVmon or EV control and 50ng p65 per well. Vpx/Vpr expression was detected using an anti-Vpx antibody. Data are mean ± SD, *n* = 3, representative of at least 3 repeats. Statistical analyses were performed using Student’s *t*-test, with Welch’s correction where appropriate. \**P* < 0.05, \*\**P* < 0.01, \*\*\**P* < 0.001.

## Discussion

In this study we evidence Vpx as an inhibitor of NF-*κ*B signaling activated by diverse agonists (Fig 1). Inhibition mapped to NF-*κ*B family member p65 (Fig 2) and was unrelated to Vpx interaction with known cellular partners including SAMHD1, HUSH complex, STING or DCAF1 (Fig 3). Vpx co-immunoprecipitated with p65, but not NF-*κ*B proteins p50 or p100 (Fig 4). Vpx did not prevent p65 phosphorylation, a marker of activation, but rather inhibited p65 nuclear translocation (Fig 5), explaining its broad activity against different NF-*κ*B activating agonists. Consistent with independence from DCAF1, which recruits the CRL4 E3 ubiquitin ligase complex to drive target protein degradation, p65 was not degraded by Vpx (Fig 5). Inhibition of p65 was found to be conserved amongst Vpx proteins from distantly related SIV, as well as HIV-2, and Vpr from SIVmon which, like Vpx, exhibits SAMHD1-degrading activity (Fig 6).

These findings extend recent observations that Vpx binds STING to suppress NF-*κ*B activation downstream of DNA sensing(28). This preceding study did not explore the role of Vpx as an NF-*κ*B signalling antagonist in the setting of cognate virus infection in the absence of pharmacological STING activation or test Vpx NF-*κ*B antagonism against the full range of NF-*κ*B agonists(28). However, similar to this work, we also found that Vpx did not inhibit cGAS/STING-induced IRF3 activation (Supplementary Fig1 E-G). However, we found that the proposed STING-binding Vpx mutant R51A S52A remained competent for NF-*κ*B antagonism downstream of STING activation. We suggest that in addition to manipulation of STING, Vpx directly targets p65 to prevent its nuclear translocation. This is the first description of a lentiviral protein targeting NF-*κ*B activity in this manner and our findings are consistent with a recent study which identified Vpx-p65 interaction using unbiased mass spectrometry (31).

We hypothesise that the relationship between lentiviruses and NF-*κ*B is complex because this transcription factor family has both anti-viral activities, e.g. downstream of DNA sensing, and pro-viral activities, as a key transcription factor in lentiviral promoters. Importantly, viral accessory proteins also have other pro-viral activities, mediated by manipulation of a further complex set of host pathways including manipulation of epigenetic regulation of transcription Vpx-HUSH(23,24), Vpr-SLF2(32), and regulation of NF-*κ*B and IRF3 nuclear transport(16). Thus, how Vpx impacts viral replication is very dependent on the nature of the assay, and the cells used. For example, during spreading infection in Jurkat T cells, which lack active SAMHD1 and cGAS, WT SIVmac and Δvpx viruses replicate similarly, unless STING is activated with agonist (RR-S2 CDA), and then Vpx enhances infection, consistent with our data (28,33). The situation is certainly complex and incompletely understood. For example, rather than having a negative effect, through inhibiting NF-*κ*B and therefore lentiviral transcription, available studies support the opposite phenomenon: Vpx degradation of HUSH enhances spreading infection of SIVmac in CEMx174 cells and HIV-1 proviral transcription in Jurkat cells with Vpx delivered by VLP (23,24). Indeed, Vpx degradation of HUSH to enhance transcription may compensate for transcription inhibition through NF-*κ*B antagonism. It has been suggested that HIV-1 Vpr may be sequestered by Gag in infected cells allowing Vpr-inhibited pathways to reactivate (34). This may also be true for Vpx, which is also incorporated into particles through Gag recruitment(35).

Classically, Vpx degradation of SAMHD1 has allowed Vpx-bearing VLP to rescue HIV-1 from SAMHD1 mediated inhibition of DNA synthesis. However, restored HIV-1 DNA synthesis cannot rescue viral replication because DNA typically activates DNA sensing by cGAS leading to induction of an antiviral state (36,37). Thus, although single round infection of macrophages is improved by Vpx, carrying Vpx does not tend to rescue lentiviral replication in DNA sensing competent cells such as macrophages, irrespective of its anti-SAMHD1 activity. On the other hand, HIV-1, which does not encode Vpx, replicates well in macrophages, waiting until cells enter a permissive G1 like state in which SAMHD1 is switched off, in order to bypass inhibition of reverse transcription (38). One possibility is that Vpx-bearing viruses use Vpx to enhance replication in T cells rather than macrophages, manipulating NF-*κ*B subtly enough to suppress anti-viral activity while retaining pro-viral transcription (39).

Pro-inflammatory cytokine secretion is activated during HIV-1 transmission *in vivo*(40). Similar data are not available for Vpx-encoding viruses, but the ability to antagonise diverse anti-pathogen signalling likely benefits transmission, particularly across mucosal surfaces where sentinel myeloid cells will limit infection if activated for example, through type 1 interferon secretion (41–43). Vpx mediated inhibition of NF-*κ*B may also benefit the virus by antagonising signalling downstream of pattern recognition receptors including TLR7 and 8: both of which detect HIV-1 RNA and activate NF-*κ*B(44).

Importantly, our study suggests an additional function for Vpx in manipulating cell biology via NF-*κ*B and this should be taken into account when using Vpx experimentally to enhance transduction of lentiviral vectors (45,46) and in *in vivo* applications, for example in lentivector gene transduction (47) and vaccine delivery (48). Furthermore, mechanistic and structural studies of Vpx targeting of p65 may inform the development of NF-*κ*B inhibitors with novel immunotherapy properties and uses in infectious and non-infectious pathologies.

## Materials and Methods

### Plasmids

Codon-optimised Vpx cDNAs were synthesised by GeneArt. and cloned into a pcDNA3.1 plasmid with or without an N-terminal FLAG tag using BamHI and NotI. Point mutations were introduced into plasmids using site directed mutagenesis with overlapping forward and reverse primers, each bearing the mutation (see Table 1). SDM PCRs were performed using Pfu Turbo DNA Polymerase (Agilent) followed by DpnI digest (NEB) according to the manufacturer’s instructions (Agilent).

**Table 1.**
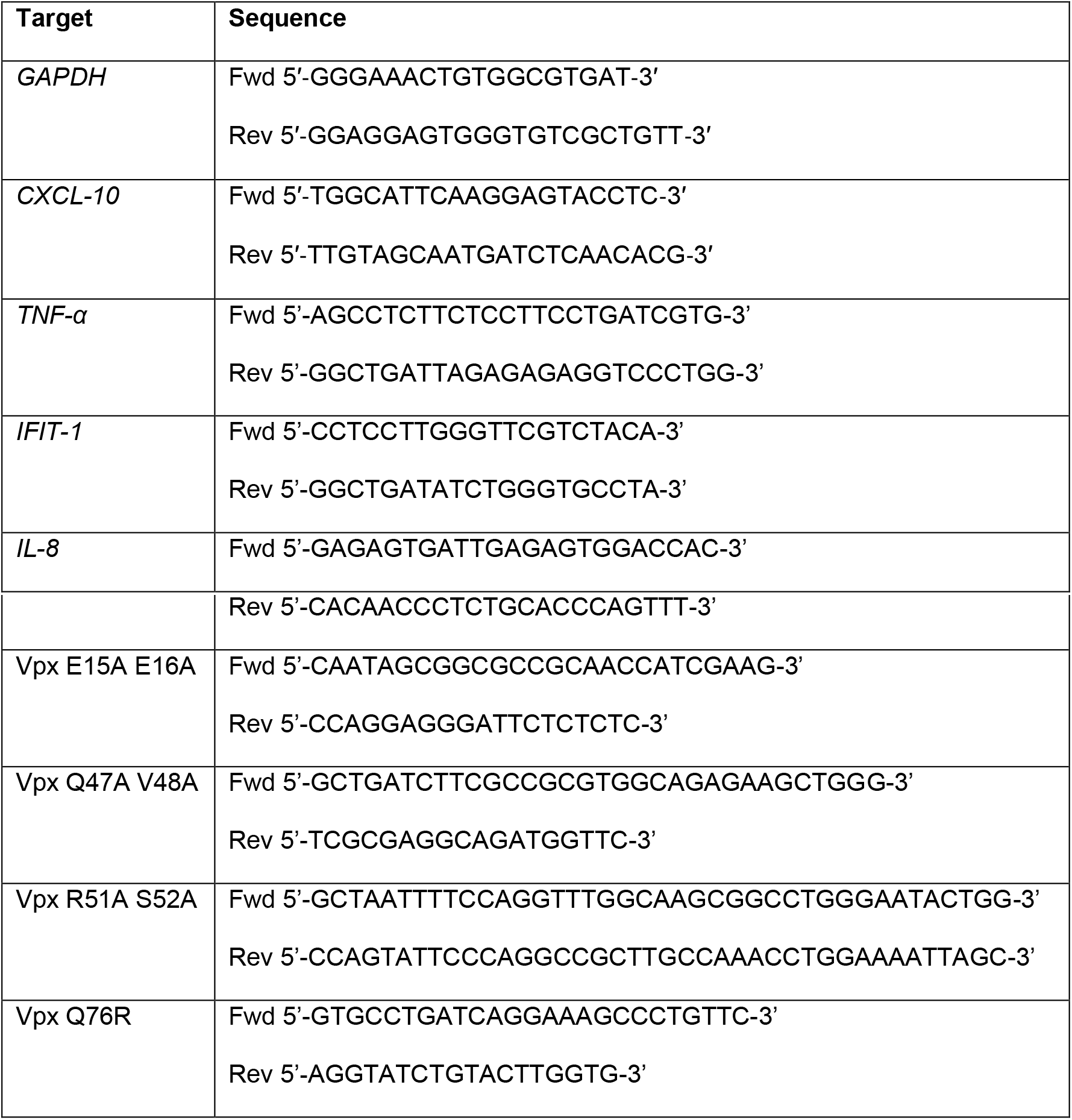
Primers.

Successful insertion of the desired mutations was confirmed by sequencing. The NF-*κ*B reporter plasmid containing five copies of an NF-*κ*B response element fused to the firefly luciferase gene and TK-renilla control plasmid were obtained from Promega. The IFIT1 (ISG56) promoter fused with firefly luciferase reporter construct was obtained from A. Bowie, Trinity College Dublin. Plasmids expressing HA-tagged p50, p65, p100 and IKKβ and FLAG-tagged cGAS and STING were generated by amplifying the relevant cDNAs and cloning them into a version of pcDNA3.1 with an N-terminal HA or FLAG tag with NotI and XbaI. Plasmids expressing TRAF2 and TRAF6 were obtained from A. Bowie, Trinity College Dublin.

### Cell lines and primary cell preparation

HEK293T and HeLa cells were grown in Dulbecco’s modified Eagle’s medium (DMEM; Gibco) supplemented with 10% fetal calf serum (FCS; Gibco) and penicillin-streptomycin (50 μg/ml) (Gibco). Peripheral blood mononuclear cells (PBMCs) were prepared from HIV seronegative donors (after informed consent was obtained), by density-gradient centrifugation (Lymphoprep, Axis-Shield, UK). Monocyte-derived macrophages (MDM) were prepared by adherence with washing of non-adherent cells after 2 h, with subsequent maintenance of adherent cells in RPMI 1640 medium supplemented with 10% human serum and M-CSF (10 ng/ml, Peprotech) for 3 days and then differentiated for a further 4 days in RPMI 1640 medium supplemented with 10% fetal calf sera without M-CSF.

### Agonists

Lipopolysaccharide (LPS), tumour necrosis factor-alpha (TNF-α) and interleukin-1 β (IL-1β) were obtained from Peprotech. Sendai virus was obtained from Charles River Laboratories.

### Transfection and small interfering RNA (siRNA) interference

For dual luciferase reporter gene assays in HEK293T cell, 1.5×10^5^ cells/ml were seeded in 24 well plates and transfected with 5ng luciferase reporter (IFIT1 or NF-*κ*B-sensitive), 2.5ng thymidine kinase renilla luciferase reporter (Promega), 0.5-200ng empty or Vpx expressing pcDNA3.1, 1.5-6ng pcDNA3.1 FLAG-cGAS and STING using 0.75μl FuGENE 6 (Promega) and 20μl Opti-MEM (Promega). All transfections were topped up with an empty vector plasmid to equalise total amounts of DNA. 48 hours later cells were lysed in passive lysis buffer (Promega), and firefly and renilla luciferase activities were measured using a Glomax luminometer (Promega). Expression of the thymidine kinase renilla luciferase reporter was used as a control for transfection efficiency between wells. The fold induction of the reporter activity was calculated by normalising each result to the luciferase activity of the unstimulated cells transfected with empty pcDNA3.1 (EV). For siRNA experimentsHEK293T cells (1.5×10^5^ cells/ml) were seeded in 6-well plates and transfected with 150nM siRNA using 4μl Lipofectamine 2000 (Invitrogen) and 184μl OptiMEM. 48 hours later 2×10^5^ cells/ml were seeded in 24-well plate to carry out the reporter gene assays.

### Virus production and infection

VLPs were produced by transfecting T150 flasks of HEK293T cells with 8μg of vesicular stomatitis virus-G glycoprotein (VSV-G) expressing plasmid pMDG (Genscript) pMDG, 32μg SIV4+ (49)and with or without 1μg of pcDNA3.1 Vpx expression plasmid using Fugene 6 transfection reagent (Promega) according to the manufacturer’s instructions. A chimeric virus derived from primary isolate SIVsm(E543) was used to investigate the function of Vpx in the context of the original SIVsm lineage(50). The chimera was made by inserting the *gag, pol* and accessory genes (*vif, vpr, vpx*) of SIVsm(E543) into an SIVmax239-based vector where a large deletion in e*nv* limits the vector to single-round infection of human cells and *GFP* is expressed in place of *nef*. SIVsm(E543) WT and Δvpx were produced by transfecting 10μg pMDG and 25μg SIVsm construct. Supernatants were harvested 48 and 72 hours post-transfection and filtered through a 0.45μm filter. All lentivectors and VLPs were DNAse treated (2 h at 37°C, DNaseI, Sigma) before they were concentrated by ultracentrifugation in a Sorvall Discovery (Hitachi) at 23000rpm for 2 hours at 4°C under vacuum conditions through a 20% sucrose cushion. The pellet was then resuspended in RPMI + 10% FCS and stored at −80°C. Viral preparations were quantified by qPCR using a SYBR Green-based product-enhanced RT (SG-PERT) assay as described to equilibrate viral dose(51). MDM were infected in the presence of 8 μg/ml polybrene (Sigma) and for SIVsm, infection levels were determined 48h later by enumerating GFP-positive cells by flow cytometry using the FACS Calibur (BD) and analysing with FlowJo software.

### Immunoblotting and immunoprecipitation

For immunoblotting cells were lysed in either passive lysis buffer (Promega) or cell lysis buffer (50 mM Tris pH 8, 150 mM NaCl, 1 mM EDTA, 10% (v/v) glycerol, 1 % (v/v) Triton X100, 0.05 % (v/v) NP-40 supplemented with protease inhibitors (Roche), and phosphatase inhibitors (Roche) for immunoblotting with phospho-specific antibodies. Lysates were clarified by centrifugation at 14,000 × *g* for 10 min and boiled in 6x protein loading buffer, containing 50 mM Tris-HCl (pH 6.8), 2 % (w/v) SDS, 10% (v/v) glycerol, 0.1% (w/v) bromophenol blue,100 mM β-mercaptoethanol for 5 min. Proteins were separated by SDS-PAGE on 12% polyacrylamide gels and transferred to a Hybond ECL membrane (Amersham biosciences) using a semi-dry transfer system (Biorad). Membranes were subsequently blocked by incubation for 1 h at room temperature in 5 % (w/v) milk proteins + 0.01 % (v/v) Tween-20 in PBS (PBST). The membranes were then incubated overnight at 4°C with primary antibody (Ab) diluted in 5 % (w/v) milk proteins in PBST. Primary antibodies were from the following sources: mouse anti-β-actin (Abcam), mouse-anti-tubulin (EMD Millipore), mouse-anti-FLAG (Sigma), rabbit-anti-HA (Sigma), mouse-anti-p65 (Santa Cruz), rabbit-anti-phospho p65 (Ser 536) (Cell Signaling), rabbit-anti-IκBα (Cell Signaling), mouse-anti-phospho-IκBα (Cell Signaling), mouse-anti-Vpx raised against HIV-2 Vpx (NIH AIDS Reagents), rabbit-anti-DCAF1 (Bethyl). Primary antibodies were detected with goat-anti-mouse/rabbit IRdye 800CW infrared dye secondary antibodies and membranes imaged using an Odyssey Infrared Imager (LI-COR Biosciences). For co-immunoprecipitation assays HEK293T cells were grown in 10 cm dishes and co-transfected with 5μg of a plasmid expressing FLAG-tagged Vpx and 5μg of a plasmid expressing HA-tagged p50, p65, p100, IKKβ or an empty vector (EV) control using polyethylenimine (Polysciences) according to the manufacturer’s instructions. After 24 h cells were lysed in lysis buffer (0.5 (v/v))% NP-40 in PBS supplemented with protease inhibitors (Roche) and phosphatase inhibitors (Roche), pre-cleared by centrifugation and incubated with 25μl of mouse-anti-HA agarose beads (Millipore) or mouse-anti-FLAG M2 agarose affinity gel (Sigma) for 2–4 hr. Immunoprecipitates were washed 3 times in 1 ml of lysis buffer and eluted from the beads by boiling in 20 μl of 2X protein loading buffer. Proteins were resolved by SDS-PAGE and detected by immunoblotting as described above.

### RNA extraction and quantitative real-time PCR (RT-PCR)

RNA was extracted using a total RNA purification kit (Norgen) according to the manufacturer’s instructions. 1μg RNA was used to synthesise cDNA using Superscript III reverse transcriptase (Invitrogen), also according to the manufacturer’s protocol. cDNA was diluted 1:5 in water and 2μl was used for real-time PCR using SYBR^®^ Green PCR master mix (Applied Biosystems) and a Quant Studio 5 real-time PCR machine (Applied Biosystems). Expression of each gene was normalised to an internal control (*GAPDH*), and these values were then normalised to mock/EV-treated control cells to yield a fold induction.

### Nuclear translocation assay

#### Image acquisition

For nuclear translocation assays, HeLa cells (5×10^4^ cells/ml) were adhered in an Cellcarrier Ultra optical 96-well plate (PerkinElmer). Cells were washed three times with ice-cold PBS and fixed in 4% (vol/vol) paraformaldehyde. The cells were permeabilised in 0.1% (vol/vol) Triton X-100 in PBS, and blocked for 1h in 10% (vol/vol) goat serum in PBS with 0.1% w/v BSA. The cells were stained with mouse-anti-p65 (Sigma) for 1 hour followed by incubation with goat anti-mouse Alexa Fluor 488 secondary IgG antibody (Life Technologies). Cells were stained with anti-flag antibody for 1h. Cells were incubated for 1h with Phalloidin-568 in PBS, washed followed by incubation for 30 minutes with 1μg/ml DAPI (4’,6-diamidino-2 phenylindole) was added per well to visualise DNA. Cells were washed with PBS three times between each step. Images were acquired using the WiScan^®^ Hermes High-Content Imaging System (IDEA Bio-Medical, Rehovot, Israel) at magnification 10X/0.4NA Four channel automated acquisition was carried out sequentially (DAPI/TRITC, GFP/Cy5). Images were acquired at 10X magnification, 100% density/ 80% well area resulting in 47 FOV/well.

#### Image analysis

NF*κ*B raw image channels were pre-processed using a batch rolling ball background correction in FIJI imagej software package prior to quantification(52). Automated image analysis was carried out using the ‘Translocation’ module of the Athena Image analysis software (IDEA Bio-Medical, Rehovot, Israel)). Firstly, Nuclei were identified as primary objects by segmentation of the Hoechst33342 channel. Cells were identified as secondary objects by nucleus depended on segmentation of the Phalloidin-A488 channel. Cell cytoplasm was segmented by subtracting the nuclear objects mask from the cell masks. Vpx positive cells were identified by identifying Vpx IF signal (FLAG-A488)signal as independent granules. Vpx+ cells were determined by the presence of Vpx+ signal within a cell border applied to filter the segmented cell population. Intensity properties were calculated for the nuclei, cytoplasm and cell object populations. Nuclear:cytoplasmic ratio (NCR) was calculated as part of the pipeline by dividing the Integrated Intensity of the nuclei object by the integrated intensity of corresponding cytoplasm object. Data post-processing was carried out in Python programming languages using the Numpy and Pandas packages. An arbitrary cut-off of NCR of less than 0.01 and greater than 10 was applied to the data to filter outliers. 10,000 cells per condition, sampled in Python were plotted using Graphpad Prism 9.

### Phylogenetic analysis

*Vpr* and *vpx* sequences obtained from the Los Alamos database were manually aligned using the Seaview sequence editor (53). Whilst *vpr* and *vpx* have undergone extensive insertions and deletions in various lineages, multiple alignment strategies consistently give the clades of interest strong support. Maximum-likelihood phylogenies were estimated using the General Time-Reversible model of nucleotide substitution with gamma-distributed rate variation across sites (GTR-gamma) implemented in RAxML 8 software(54). Branch support was determined using 1000 bootstrap alignments. Phylogenies were visualized with the program FigTree (https://github.com/rambaut/figtree/releases). The vpr/vpx tree was midpoint-rooted. Branch lengths indicate the number of nucleotide substitutions per site.

### Statistical analysis

Statistical analyses were performed using an unpaired Student’s *t*-test with Welch’s correction where variances were unequal. \**P* < 0.05, \*\**P* < 0.01, \*\*\**P* < 0.001, **** *P* < 0.0001.

## Acknowledgements

GJT was funded through a Wellcome Trust Senior Biomedical Research Fellowship (108183) followed by a Wellcome Investigator Award (220863), the European Research Council under the European Union’s Seventh Framework Programme (FP7/2007-2013)/ERC (grant HIVInnate 339223) a Wellcome Trust Collaborative award (214344) and the National Institute for Health Research University College London Hospitals Biomedical Research Centre. DLF was funded by a Wellcome Research Training Fellowship (174356). CMM was funded by the BBSRC (BB/T006501/1). JC was supported by the Wolfson Foundation.

