## Supplementary figures for "HIV-2/SIV Vpx antagonises NF-*κ*B activation by targeting p65"

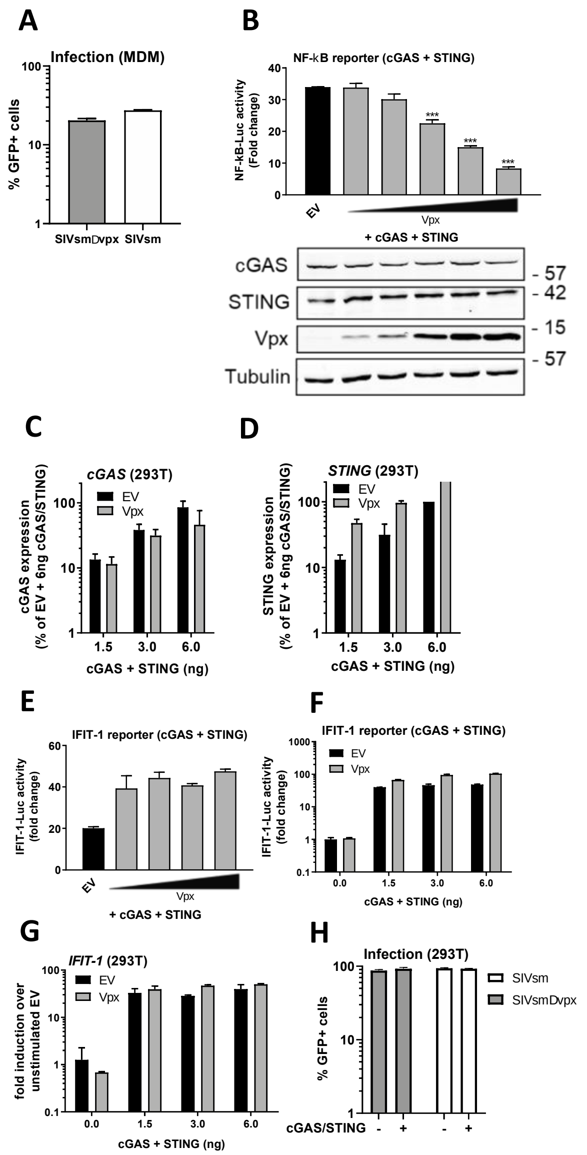


Supplementary Figure 1. Vpx is a broad antagonist of NF-$\kappa$B

A: Infection data from Fig 1A and B. MDM infected for 48h with SIVsm or SIVsmΔVpx expressing GFP (1.5 U/ml RT).

B: NF-$\kappa$B reporter activity and immunoblot from HEK293T cells co-transfected for 24h with 1-100ng SIVmac Vpx or EV control (100ng) and 1.5ng each of FLAG-cGAS and STING per well. cGAS and STING were detected using a FLAG antibody.

C: *cGAS* qRT-PCR from HEK293T cells co-transfected for 24h with 50ng SIVmac Vpx or EV control plus 1.5, 3 or 6ng each of FLAG-cGAS and FLAG-STING per well.

D: *STING* qRT-PCR from HEK293T cells co-transfected for 24h with 50ng SIVmac Vpx or EV control plus 1.5, 3 or 6ng each of FLAG-cGAS and FLAG-STING per well.

E: IFIT-1 reporter activity from HEK293T cells co-transfected for 24h with 12.5-100ng SIVmac Vpx or EV control (100ng) and 1.5ng each of FLAG-cGAS and STING per well.

F: IFIT-1 reporter activity from HEK293T cells co-transfected for 24h with 50ng SIVmac Vpx or EV control plus 0, 1.5, 3 or 6ng each of FLAG-cGAS and FLAG-STING per well.

G: *IFIT-1* qRT-PCR from HEK293T cells co-transfected for 24h with 50ng SIVmac Vpx or EV control plus 0, 1.5, 3 or 6ng each of FLAG-cGAS and FLAG-STING per well.

H: Infection data from Fig 1I. HEK293T infected for 48h with SIVsm or SIVsmΔVpx expressing GFP (1.0 U/ml RT).

Data are mean ± SD, $n = 3$, representative of at least 3 repeats. Statistical analyses were performed using Student's *t*-test, with Welch's correction where appropriate. ***$P < 0.001$.

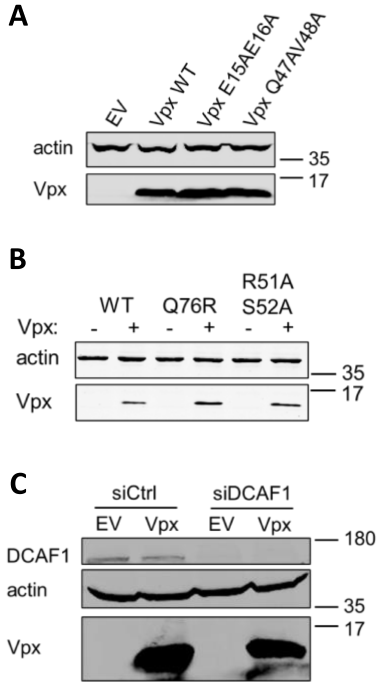


Supplementary Figure 2. Vpx-mediated inhibition of NF-κB is independent of DCAF1, SAMHD1 and HUSH

A: Immunoblot from NF-κB reporter activity in HEK293T cells from Fig 3B detecting Vpx mutants using an antibody against the FLAG tag and actin.

B: Immunoblot from NF-κB reporter activity in HEK293T cells from Fig 3C, D detecting Vpx mutants using an antibody against the FLAG tag and actin. The highest transfection dose was selected.

C: Immunoblot for DCAF1 depletion in NF-κB reporter activity in HEK293T cells from Fig 3E, F detecting DCAF1, actin and Vpx using a Vpx antibody.

Data are from a representative experiment repeated at least 3 times.

**A**

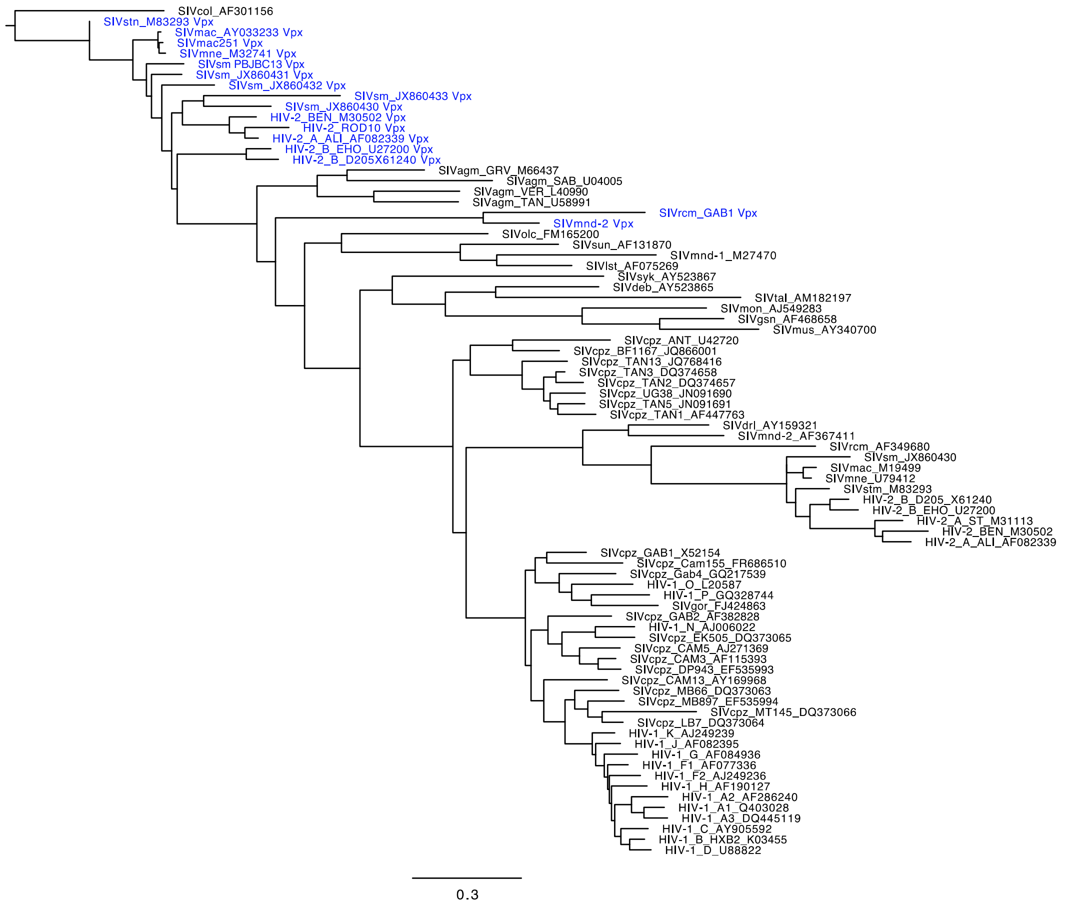


Supplementary Figure 6. Inhibition of NF-$\kappa$B is conserved amongst Vpx species variants

A. Maximum likelihood phylogeny of primate lentiviruses *vpr* and *vpx* genes generated using FigTree software. Vpx sequences are coloured blue; vpr sequences are coloured black. Horizontal branch lengths are shown to scale. Scale bar represents 0.3 nt substitutions per site.
